# How host self-regulation governs hyperparasitoid persistence in a caterpillar-parasitoid-trigonalid system

**DOI:** 10.64898/2026.09.28.755095

**Authors:** Zyanya R. Rosales, Josué Manik Nava-Sedeño

## Abstract

Species at the highest trophic levels are generally the least resilient to perturbations, yet the factors that determine their persistence remain poorly understood. We ask how the intrinsic self-regulation of a basal host governs the persistence of the trophic levels above it. To address this question, we use a minimal discrete-time model of a host–parasitoid–hyperparasitoid system motivated by the biology of trigonalid wasps, obligate hyperparasitoids that complete development only when their caterpillar host is also attacked by a primary parasitoid. We compare two forms of host self-regulation in the absence of natural enemies: scramble (Ricker) fecundity, appropriate when the parasitoid is the host’s principal enemy and which recovers the classical model of Beddington and Hammond as a special case; and contest (Beverton–Holt) fecundity, appropriate when the host is held in check by many additional enemies. Using numerical continuation, Lyapunov exponents, and direct computation of the model’s trajectories, we find that the form of host regulation is decisive for persistence at the top trophic level. Under scramble host dynamics, coexistence is extremely fragile with respect to both parameter values and initial conditions. Under contest host dynamics, coexistence is robust across broad parameter ranges and loses stability only gradually, through a supercritical Neimark–Sacker bifurcation and a subsequent torus-doubling route to chaos. Therefore, in at least some trophic networks, the resilience of the highest levels can be determined at the base of the network.

## 1 Introduction

The dynamics of ecological populations are among the earliest problems studied in mathematical biology. Initial work focused on self-regulated single populations and was soon extended to interactions among populations. The accelerating extinction of species and the increasing fragility of present-day ecosystems have motivated a plethora of mathematical models that investigate the mechanisms underlying biodiversity loss and explore possible future scenarios [1–5]. A particularly important topic in this context is population resilience [6], i.e. the maintenance of a population’s longterm dynamics despite changes in population size (initial conditions) or behavioral and environmental conditions (parameter values). In particular, populations at the highest trophic levels have been shown to be much less resilient than those at lower levels [7–9].

In invertebrate trophic networks, such as those of terrestrial arthropods, parasitic species use hosts (usually insect larvae) as food for their own larvae, killing the host in the process. Hyperparasitic species, in turn, parasitize primary parasitoids in the same manner. Thus, hyperparasitoids occupy some of the highest levels of the trophic network. Among the most biologically interesting hyperparasitoids are wasps of the family *Trigonalidae* [10–12]. These wasps lay their eggs along the edges of leaves eaten by butterfly or moth larvae, i.e. caterpillars. A caterpillar feeding on a leaf bearing trigonalid eggs may accidentally ingest them. The caterpillar’s mandibular movements and digestive fluids trigger the emergence of trigonalid larvae from the eggs. A trigonalid larva then bores into the caterpillar’s haemocoel, or circulatory system, where it remains dormant without harming the caterpillar. If more than one trigonalid larva is present in the same host, they typically compete until only one survives. If the caterpillar is subsequently parasitized by a primary parasitoid wasp, for example one from the family *Ichneumonidae*, the trigonalid larva parasitizes the developing ichneumonid larva. In this scenario, both the caterpillar and the ichneumonid larva die, while the trigonalid larva successfully develops into an adult wasp. If the caterpillar is never parasitized by an ichneumonid wasp, the trigonalid larva fails to develop and dies within the caterpillar’s body. Some trigonalid species, such as *Taeniogonalos venatoria* and *Taeniogonalos maculata*, can act as facultative primary parasitoids; however, these species parasitize sawflies of the family *Pergidae* rather than caterpillars [10, 13]. Trigonalid wasps are rarely observed, although the family itself is highly diverse.

Although host–parasitoid–hyperparasitoid systems have been studied using mathematical models [14–16], most previous studies focus heavily on the parameter regimes required for the stability of a coexistence steady state, the effects of introducing a hyperparasitoid into a host–parasitoid system, and the dynamical effects of varying a particular biological quantity. Furthermore, these models generally treat the host as a pest to be controlled by the parasitoid. As far as we are aware, the ecological resilience of species in a host–parasitoid–hyperparasitoid system has not been studied. In particular, previous studies have not focused on the resilience of the highest trophic level, the hyperparasitoid.

In this work, we study whether the form of the host’s intrinsic self-regulation, i.e. the dynamics that the host would follow in the absence of both the parasitoid and the hyperparasitoid, determines the survival of the trophic levels above it. We address this question by comparing two host fecundity functions that represent different forms of intraspecific dynamics and correspond to different ecological settings: Ricker and Beverton–Holt functions. We first consider a Ricker-type fecundity function, which corresponds to scramble competition among hosts at high density [17]. These dynamics are natural when the primary parasitoid is the host’s principal enemy. Removing the parasitoid then leaves the host without external population control, so the host regulates itself only at the high densities at which intraspecific competition becomes important. In this case, our system reduces to the host-parasitoid-hyperparasitoid model in [14]. The reduction is formal rather than biological. In their model, the hyperparasitoid searches for and attacks hosts that have already been parasitized, so that hyperparasitoid recruitment follows from a sequence of two attacks. Trigonalids, by contrast, never search for a host: their eggs are scattered over the vegetation and are ingested by hosts irrespective of whether those hosts are parasitized, and recruitment requires the joint occurrence of two events, namely ingestion of an egg and subsequent parasitism by a primary parasitoid. Because we assume these two events to be independent, the recruitment term factorizes into a product of encounter probabilities that is formally identical to the sequential-attack structure of [14]. Two distinct biological mechanisms therefore converge top the same model. This extends the applicability of their results to a life history for which they were not derived, and it means that the Beverton–Holt scenario studied below inherits the same generality. The analysis in [14] was restricted to characterizing the region of parameter space containing stable equilibria or simple oscillations, because the numerical tools needed to reveal the complete bifurcation structure of difference equations were still in their early stages. However, the authors were familiar with the model’s complex dynamical behavior, since an earlier study [18] first reported chaotic dynamics in a host–parasitoid model. Using several numerical approaches, we show that this system possesses a rich dynamical structure that ultimately determines whether the hyperparasitoid survives in the long term. We find that the coexistence steady state is fragile with respect to both parameter changes, which represent environmental or behavioral changes, and changes in the initial conditions, which represent fluctuations in population size. For most parameter values and initial conditions, the hyperparasitoid population quickly crashes, after which the dynamics continue in the two-dimensional host–parasitoid phase space. This fragility results from transcritical and Neimark–Sacker bifurcations for slow host reproduction and from flip and Neimark–Sacker bifurcations for fast host reproduction. The system also exhibits dangerous, extreme oscillations that leave all three populations susceptible to extinction under random fluctuations. These oscillations arise from bifurcations of the periodic and quasiperiodic attractors that appear when the coexistence state becomes unstable.

When enemies other than the parasitoid also prey on the host, they prevent the host population from reaching the high densities needed to trigger its intrinsic scramble dynamics, even in the absence of the primary parasitoid. We therefore also consider a Beverton–Holt fecundity function for the host, as discussed in [19]. In contrast to the previous scenario, this system exhibits much simpler dynamics. We did not observe chaotic dynamics in the hyperparasitoid-free system, and in the full system the coexistence equilibrium is resilient to changes in both initial conditions and parameter values. Even when the coexistence equilibrium becomes unstable and an invariant closed curve appears, the new attractor is itself extremely resilient and disappears in favor of a chaotic attractor only at extremely high parameter values.

Thus, our model shows that the dynamics of lower trophic levels strongly affect the survival of upper trophic levels. Tight control of the host through several predation pathways is essential for hyperparasitoid survival, even though high host abundance and rapid host growth are crucial for the reproduction of both the parasitoid and the hyperparasitoid. Therefore, the highest trophic levels may not be intrinsically less resilient; instead, their resilience may depend on the population control exerted at lower trophic levels.

## 2 Model definition

Because trigonalid wasps and common primary parasitoid species have short life spans, we model their population dynamics as a discrete dynamical system. We do not explicitly model the dynamics of the vegetation on which the host species feeds. Instead, we assume that, in the absence of parasitism, host growth is bounded by resource limitations and/or external predation, as specified below. Our model is based on the following assumptions:

- Surviving hosts proliferate according to growth-limited dynamics. In the absence of parasitoids, the number of hosts in generation *n* + 1, *X*_*n*+1_, depends on the number in generation *n, X*_*n*_, according to

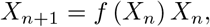

where *f* (*X*_*n*_) is the net *per capita* number of descendants produced by hosts in generation *n*. Note that *f* (*X*_*n*_) depends on the pre-parasitism population, since caterpillars compete for vegetation at the larval stage.
- Encounters between hosts and primary parasitoids, and between hosts and hyperparasitoid eggs, are modeled as Poisson-distributed events, following [20].
- A single encounter between a host and a hyperparasitoid egg results in the implantation of a hyperparasitoid larva. The number of encounters depends on the number of hyperparasitoids, *Z*_*n*_, and the average number of eggs laid per hyperparasitoid, *N*. Thus, the number of hosts carrying a hyperparasitoid larva in generation *n* is given by

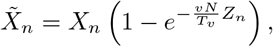

where *v* is the vegetation biomass covered per hyperparasitoid egg and *T*_*v*_ is the total vegetation biomass in the area occupied by the host.
- If a host incubating a hyperparasitoid larva does not encounter a primary parasitoid, the host develops and reproduces normally while the hyperparasitoid larva perishes; i.e. the hyperparasitoid does not directly harm the host.
- A single encounter between a host and a primary parasitoid from population *Y*_*n*_ results in the death of the host and the production of a new generation of parasitoids. Thus, the number of hosts surviving parasitism depends only on the number of primary parasitoids and is given by

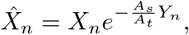

where *A*_*s*_ is the area searched per primary parasitoid and *A*_*t*_ is the total area occupied by the host.
- An encounter between a primary parasitoid and a host carrying a hyperparasitoid larva results in the deaths of the primary parasitoid larva and the host, as well as the successful development of the hyperparasitoid larva. Therefore, the number of primary parasitoids in generation *n* + 1 is given by

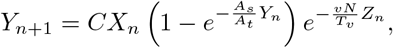

while the corresponding number of hyperparasitoids is given by

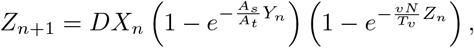

where *C* and *D* are the primary parasitoid and hyperparasitoid brood sizes, respectively.

Under these assumptions, our model in nondimensional form is given by the following system of nonlinear difference equations:

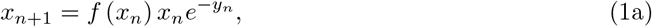

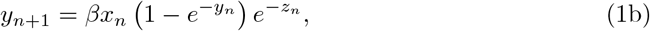

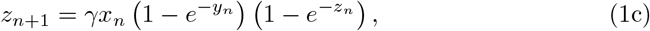

where *β* and *γ* are nondimensional parameters, given by

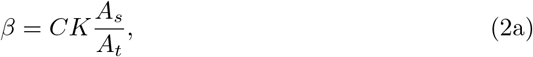

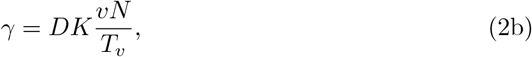

where *K* is an upper bound on the carrying capacity of the host population, in the absence of parasitoids. Thus, the nondimensional parameters are always nonnegative. The nondimensional populations in generation *n* are given by

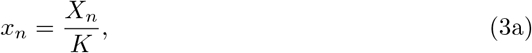

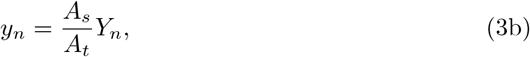

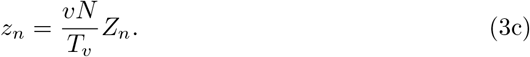

We consider two fecundity functions that model different ecological scenarios in the absence of parasitoids: a Ricker-type function, corresponding to the case in which the parasitoid is the host’s only enemy, and a Beverton–Holt-type function, corresponding to the case in which enemies other than the parasitoid also regulate the host.

### 2.1 Ricker-type fecundity

Assuming that, in the absence of parasitoids, intraspecific competition for vegetation is the primary mechanism regulating host population growth, we propose the following nondimensional Ricker-type fecundity function:

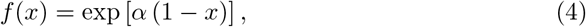

where exp(*α*) is the fecundity of the host population in the exponential-growth regime, *x≈* 0. This formulation models a scenario in which enemies other than the primary parasitoid are largely absent. For this type of fecundity, *K* represents the carrying capacity of the host population. Note that, in this case, the greater the value of *α*, the more prolific the host.

### 2.2 Beverton–Holt-type fecundity

The Ricker formulation assumes that the primary parasitoid is the host’s main source of predation. However, the hosts considered here are primarily caterpillars, which have high energy content and a high protein-to-mass ratio. Moreover, the three species considered here do not normally live in isolation. We therefore assume that, in the absence of parasitism, the host population would not reach the high levels predicted by Ricker dynamics, because other enemies, such as birds, reptiles, mammals, and arthropods, would also regulate it. We thus propose [19] the following nondimensional Beverton–Holt-type fecundity function:

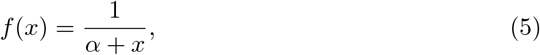

where the nondimensional parameter *α* is given by

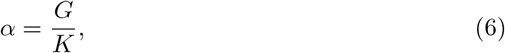

which is the ratio of the population size *G*, at which offspring production equals half the asymptotic maximum recruitment, to the asymptotic maximum recruitment *K*. For the host to survive in the absence of parasitoids, *G < K*, and therefore *α <* 1. Note that, in this case, the lower the value of *α*, the more prolific the host.

## 3 Results

The hyperparasitoid-free system, *z*_*n*_ = 0, is invariant. Its attractors therefore remain on the *xy* plane of the three-dimensional phase space, and their basins of attraction compete with those of attractors outside that plane. Thus, we study both Ricker- and Beverton–Holt-type models with and without hyperparasitoids. Because of the biological meaning of the dynamical variables, we restrict the analysis to the first octant of phase space. We study the extinction equilibria analytically and analyze the coexistence state numerically using MatContM [21].

### 3.1 Ricker-type fecundity

In the hyperparasitoid-free scenario, the system reduces to

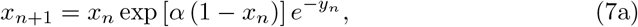

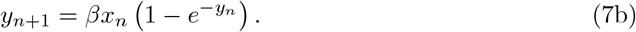

This is a nondimensional version of the model proposed in [18]. Their findings show that, for low values of *α* and *β*, the coexistence state is asymptotically stable, so the host and parasitoid populations approach constant values in the long term. As both parameters increase, this state undergoes a supercritical Neimark–Sacker bifurcation; the populations then cease to approach constant values and instead oscillate smoothly across successive generations. Further increases in both parameters induce resonance and create a stable period-5 cycle, producing dramatic population oscillations that complete a cycle every five generations. At still higher parameter values, a stable strange attractor appears, and the populations oscillate dramatically and irregularly from one generation to the next. Because oscillations may produce low population levels that are susceptible to environmental fluctuations or human impacts, the populations are resilient only at low values of *α* and *β*, i.e. when the host has low fecundity in isolation, each infected host produces few parasitoids, and/or the parasitoid has low search efficiency.

Before analyzing the full bifurcation structure, we verified that our system reduces to the system in [14]. The full system, Eqs. 1 with the Ricker-type fecundity function in Eq. 4, generally has five equilibria: the total-extinction state, *x*_*n*_ = *y*_*n*_ = *z*_*n*_ = 0; the hyperparasitoid-free state, *x*_*n*_, *≠ y*_*n*_ 0 and *z*_*n*_ = 0; the parasitoid- and hyperparasitoid-free state, *x*_*n*_ ≠ 0 and *y*_*n*_ = *z*_*n*_ = 0; and two distinct coexistence states. The two coexistence equilibria exist and are non-negative only at intermediate values of *β* and sufficiently high values of either *α* or *γ*. Thus, host fecundity in isolation, the number of primary parasitoids and hyperparasitoids produced per infected host, or hyperparasitoid egg production and dispersal must be sufficiently high for the hyperparasitoid to survive, whereas primary parasitoid efficiency must remain tightly constrained. As *β* decreases, a transcritical bifurcation first occurs between a coexistence equilibrium and the hyperparasitoid-free equilibrium, causing the latter to become a saddle. A second transcritical bifurcation then occurs between the hyperparasitoid-free saddle and the remaining coexistence equilibrium. These bifurcations indicate the existence of a critical threshold in primary parasitoid efficiency below which the hyperparasitoid cannot survive.

To study the robustness of the populations against fluctuations in population size, we estimated the basin of attraction computationally. We considered a grid of 200 *×* 200 *×* 200 initial conditions in phase space, in the interval (0, 2)^3^ *⊂* R^3^. The phase space points were classified as “converging” to the equilibrium if, after 150 steps, the trajectories were an Eculidean distance away from the equilibrium of less than 0.01, and “not converging” otherwise. We plotted the boundaries between these two regions. For low values of *α*, high values of *γ*, and appropriate values of *β*, the stable coexistence equilibrium has a small but well-defined basin of attraction, and orbits from most nearby initial conditions converge to it (Fig. 1a). However, at these low values of *α*, the parameter windows in which the coexistence equilibrium lies in the first octant and remains stable are extremely narrow: approximately one unit in *β* and 0.8 units in *γ* (Figs. 2a and c). The equilibrium is therefore highly susceptible to changes in the primary parasitoid’s behavior.

**Fig 1:**
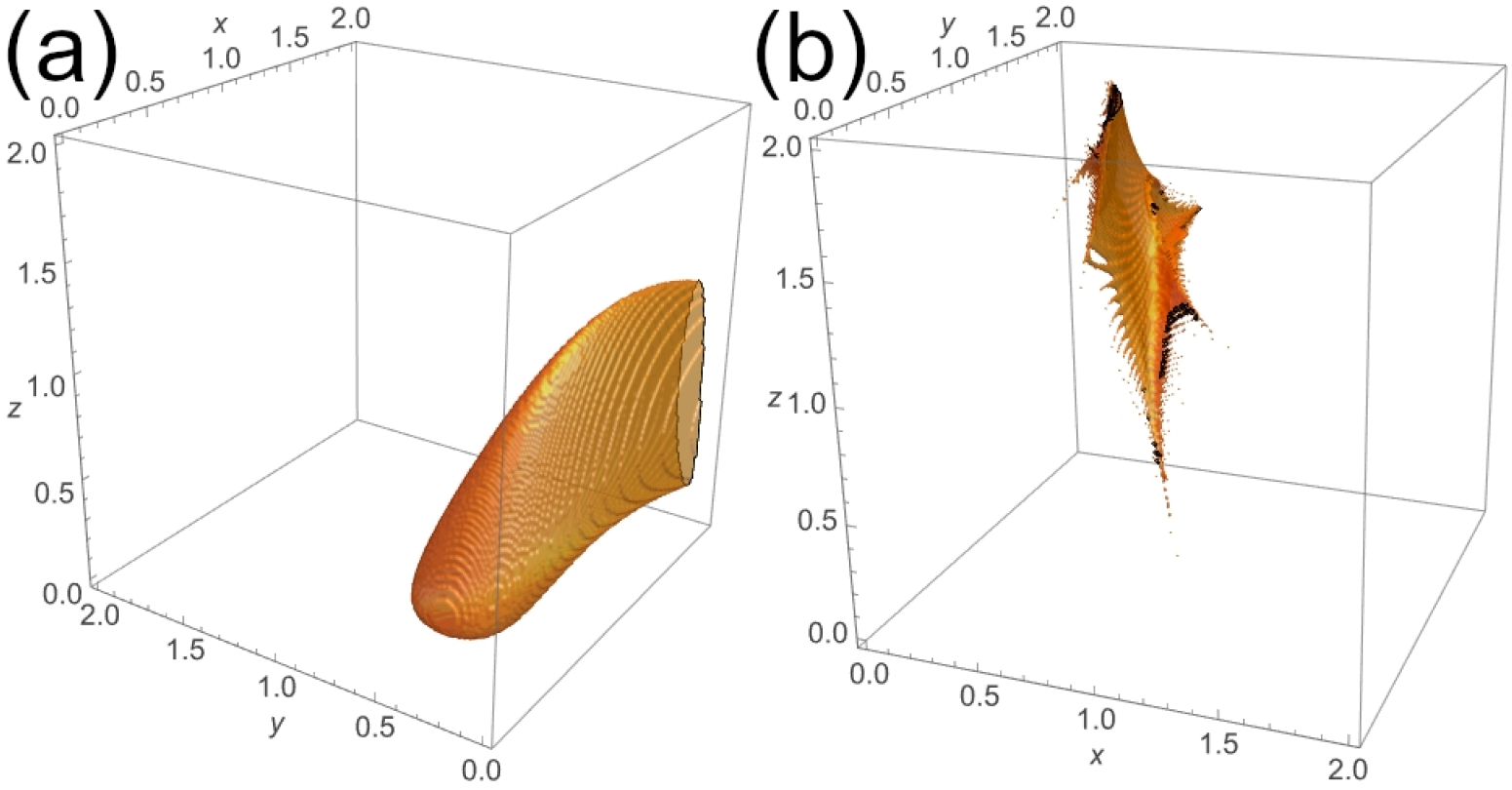
Approximate basins of attraction of the coexistence steady state for the model with Ricker-type host fecundity. Parameter values: (a) *α* = 0.85, *β* = 2.7, *γ* = 6; (b) *α* = 4, *β* = 10, *γ* = 4.

**Fig 2:**
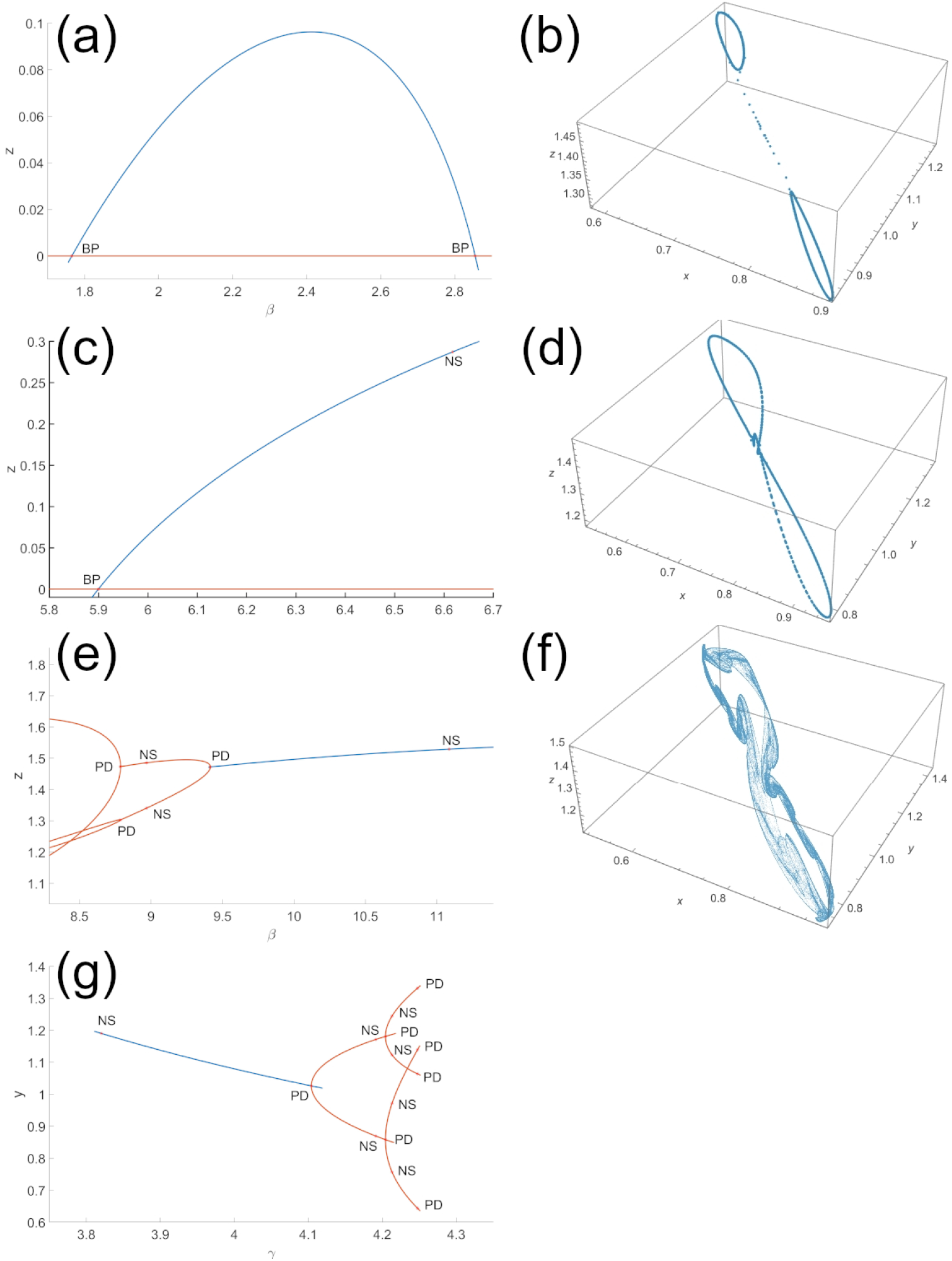
Bifurcations of the model with Ricker-type host fecundity. The left column shows local bifurcations of the coexistence steady state, while the right column shows global bifurcations emerging from the period-2 cycle. Parameter values: (a) *α* = 0.85, *γ* = 6; (b) *α* = *γ* = 4, *β* = 8.9, (c) *α* = 0.85, *β* = 2.7; (d) *α* = *γ* = 4, *β* = 8.7; (e) *α* = *γ* = 4; (f) *α* = *γ* = 4, *β* = 8.596; (g) *α* = 4, *β* = 10.

At higher values of *α*, the range of *β* values for which the coexistence state lies in the first octant remain qualitatively unchanged. However, two additional complications arise. First, invariant curves, periodic cycles, and strange attractors occupy a substantial portion of the *xy* plane, and their large basins of attraction compete with that of the coexistence equilibrium. Consequently, the basin of attraction of the coexistence state becomes smaller and more difficult to define numerically (Fig. 1b). Most initial conditions then lead to hyperparasitoid extinction, after which the system converges to the hyperparasitoid-free dynamics described above. Second, although the coexistence equilibrium remains in the first octant at high *α*, changes in either *β* or *γ* cause it to undergo a supercritical Neimark–Sacker bifurcation or a supercritical flip bifurcation, depending on the parameter values. Thus, the coexistence state is again stable only within a very narrow parameter range (Figs. 2e and g), while also having a reduced basin of attraction. The complex behavior of the system may be related to a numerically detected codimension-2 flip–Neimark–Sacker bifurcation involving *α* and *β*. This bifurcation occurs near *x ≈* 0.799, *y ≈* 0.642, *z ≈* 1.775, *α ≈* 3.202, *β* = 10, and *γ ≈*5.640, where the eigenvalues of the steady state are *λ*_1_ = *−*1 and *λ*_2,3_ *≈* exp 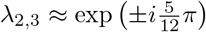. The rich dynamics are also consistent with a 1:2 resonance bifurcation involving *α* and *β*, found near *x ≈* 0.710, *y ≈* 1.396, *z ≈* 1.342, *α ≈* 4.817, *β* = 10, and *γ* ≈ 3.400, with *λ*_1,2_ =*−* 1 and *λ*_3_*≈*0.516.

Beyond the flip bifurcation, further parameter changes cause the period-2 orbit to undergo a Neimark–Sacker bifurcation, forming two invariant closed curves around its two unstable points (Fig. 2b). These curves then undergo a global bifurcation and merge into a single invariant curve that winds around both points of the unstable period-2 cycle (Fig. 2d). The merged curve subsequently undergoes resonance, producing a long-period orbit (not shown) and, ultimately, a stable strange attractor (Fig. 2f).

This analysis reveals the model’s rich dynamical behavior. Biologically, however, this complexity restricts the parameter values and initial conditions that permit coexistence, making the coexistence of all three populations fragile.

### 3.2 Beverton–Holt-type fecundity

First, we consider the hyperparasitoid-free system

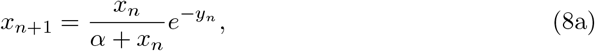

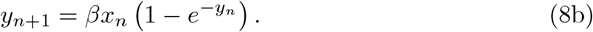

The system has three equilibria: the total-extinction state, the parasitoid-free state, and the coexistence state. A straightforward stability analysis shows that the two extinction states undergo a transcritical bifurcation at *α* = 1, where the nontrivial parasitoid-free state leaves the biologically relevant first quadrant of phase space. For *α <* 1, the parasitoid-free and coexistence states undergo another transcritical bifurcation at *β* = 1*/*(1 *−α*). At this point, the coexistence state moves from the fourth to the first quadrant and changes from unstable to stable. Further increases in *β* cause the coexistence state to undergo a supercritical Neimark–Sacker bifurcation (Figs. 3a and b), at which it becomes unstable and produces a stable invariant closed curve. As *β* increases further, the invariant curve repeatedly fragments into periodic orbits and reforms through mode locking. We confirmed this behavior numerically by calculating the average rotation number [22] of the orbit after transients, relative to the unstable coexistence equilibrium (*x*^*∗*^, *y*^*∗*^), which we treated as the center of the invariant curve. Defining

**Fig 3:**
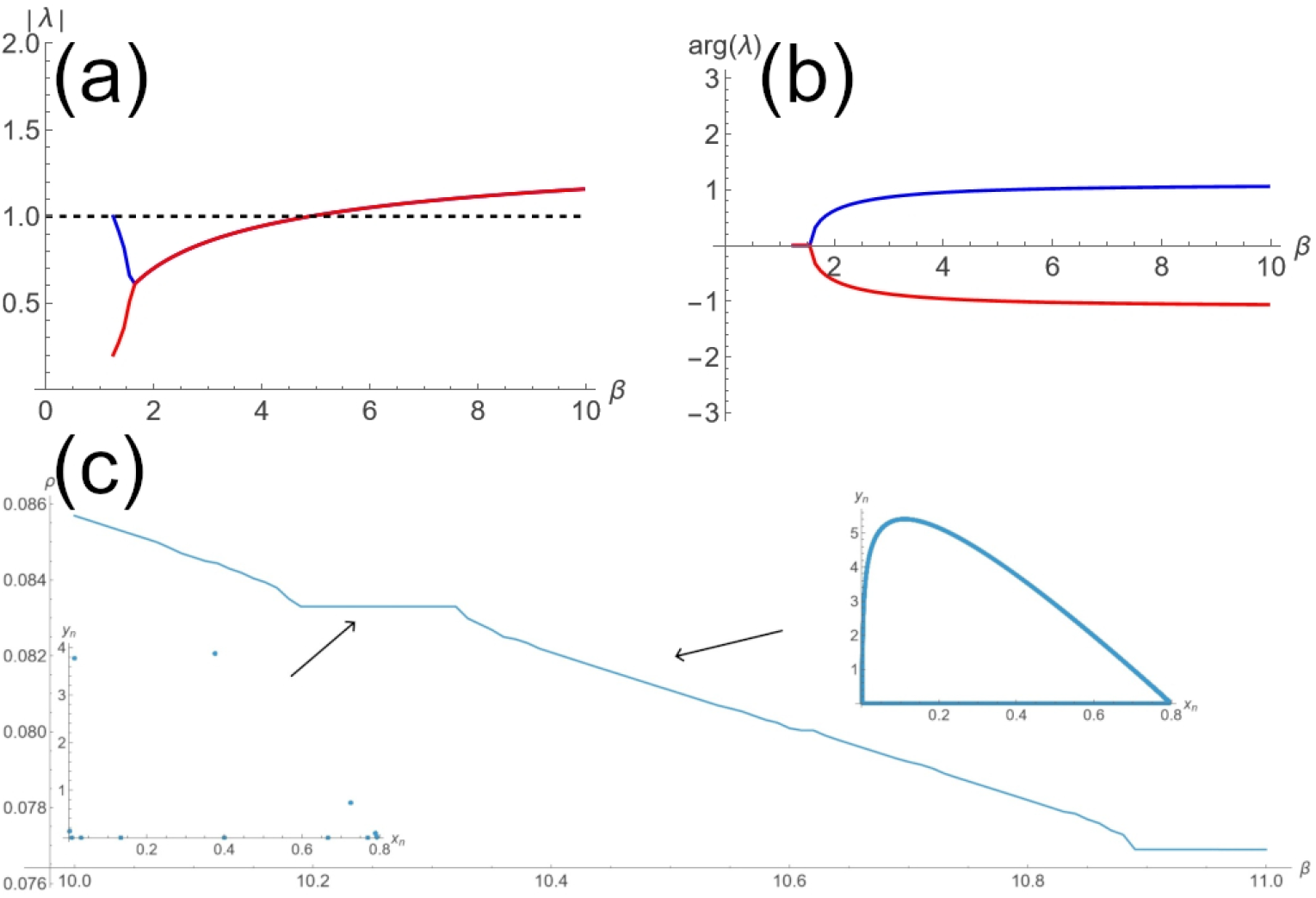
Dynamics of the hyperparasitoid-free system with Beverton–Holt host fecundity. Top: the complex-conjugate eigenvalues cross the unit circle, confirming a Neimark–Sacker bifurcation, with the remaining parameter fixed at *α* = 0.2. Bottom: the average winding number after the resulting invariant curve appears, including regions of phase locking as *β* varies and characteristic post-transient orbits. The arrows indicate the regions of the plot corresponding to the orbit in each inset. Left inset: *α* = 0.2, *β* = 10.25. Right inset: *α* = 0.2, *β* = 10.46.

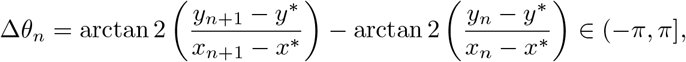

we approximated the average winding number as

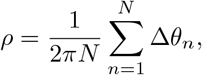

averaging over *N* = 1 *×*10^4^ orbit points after discarding the same number of transient steps. A plot of the numerical average rotation number *ρ* against *β* (Fig. 3c) exhibits a characteristic devil’s-staircase-like structure [23]. Plateaus correspond to mode-locked periodic orbits, whereas quasiperiodic orbits are more likely to occur over intervals in which *ρ* appears to decrease [24]. Importantly, we observed no chaotic attractor in the *xy* plane for any of the parameter values explored.

The full system, Eqs. 1 with the Beverton–Holt-type fecundity function in Eq. 5, generally has the same five equilibria as the system with Ricker-type fecundity. For *α≈* 1 and low values of *γ*, only hyperparasitoid-free states exist in the first octant. At sufficiently low values of *α* and/or high values of *γ*, coexistence states exist and are nonnegative. The requirement of small *α* for coexistence corresponds to rapid host growth, i.e. a small host population producing many offspring relative to the host carrying capacity. Conversely, high *γ* can correspond to a large host carrying capacity, extensive vegetation coverage by hyperparasitoid eggs, high egg production by the hyperparasitoid, and/or a small vegetation biomass occupied by the host. As before, we estimated the basins of attraction, but with a 100 *×* 100 *×* 100 grid in the region (0, 1)^3^. When the host is prolific (small *α*), the coexistence equilibrium is much more resilient to both fluctuations in population size, represented by changes in the initial conditions (Fig. 4a), and environmental or behavioral changes, represented by changes in parameter values (Figs. 4c and e). For more slowly reproducing hosts, the coexistence equilibrium is generally less resilient to fluctuations in population size (Fig. 4b), although it remains robust to environmental and behavioral changes (Figs. 4d and f). Even in this case, the equilibrium is substantially more resilient than under Ricker-type host dynamics. For any value of *α*, sufficiently large increases in *β* or *γ* cause a supercritical Neimark–Sacker bifurcation that produces a stable invariant closed curve. Under Beverton–Holt host dynamics, the bifurcations of the coexistence equilibrium are consistently simple and qualitatively similar, in contrast to the more complicated bifurcations observed under Ricker host dynamics.

**Fig 4:**
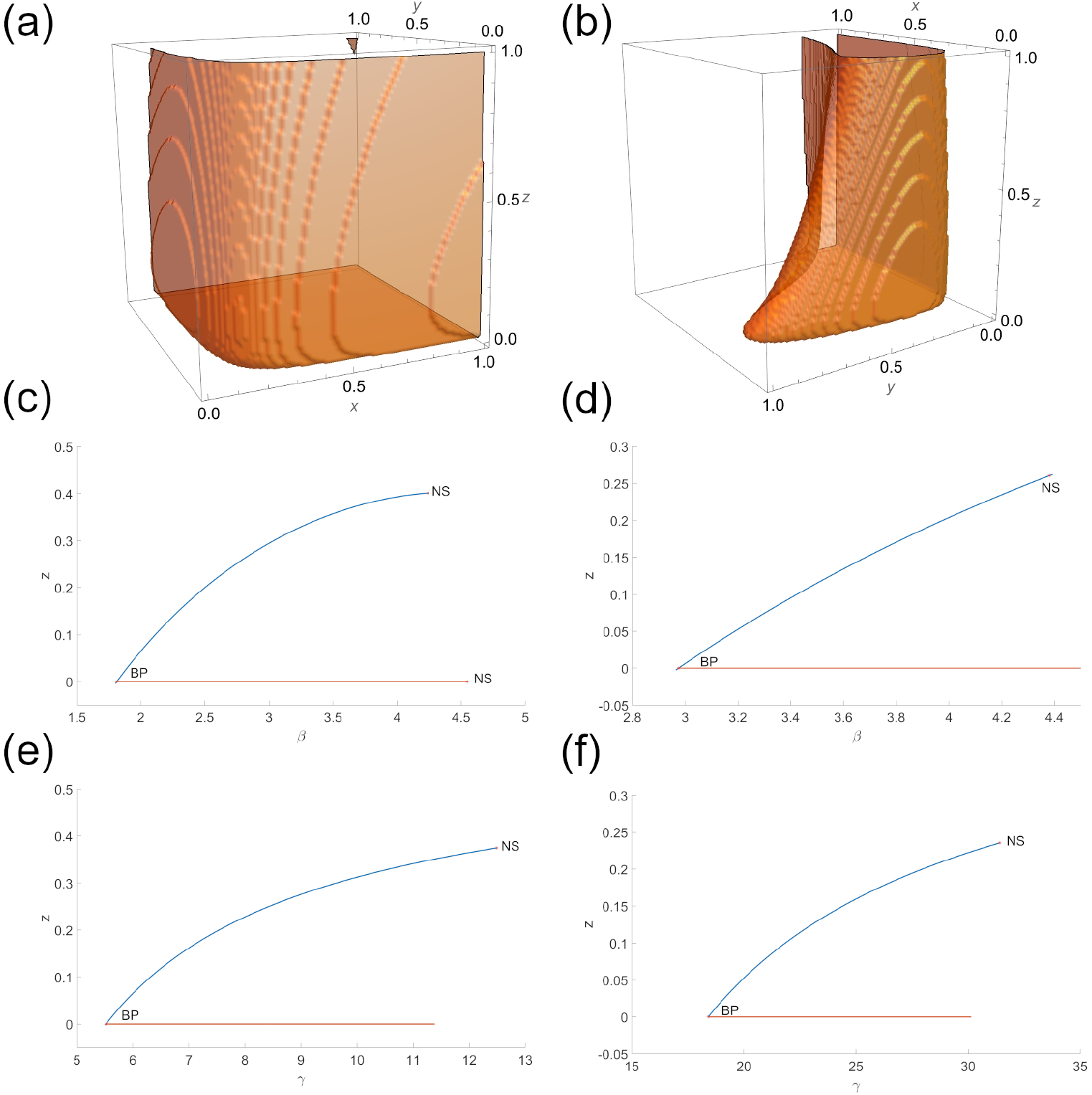
Dynamics near the coexistence state of the model with Beverton–Holt host fecundity for prolific (small *α*, left column) and slowly reproducing (large *α*, right column) hosts. Top row: numerical approximations of the basins of attraction. Middle row: bifurcation diagrams changing *β*. Bottom row: bifurcation diagrams changing *γ*. Parameter values: (a) *α* = 0.1, *β* = 2, *γ* = 6; (b) *α* = 0.5, *β* = 3.2, *γ* = 20; (c) *α* = 0.1, *γ* = 6; (d) *α* = 0.5, *γ* = 20; (e) *α* = 0.1, *β* = 2; (f) *α* = 0.5, *β* = 3.2.

Interestingly, in contrast to the system with Ricker host fecundity, the oscillations that arise after the equilibrium (Fig. 5a) becomes unstable remain small in amplitude and regular over a broad range of parameter values. Only large increases in either *β* or *γ* cause successive doublings in the length of the invariant curve (Figs. 5b– d). This behavior persists across a substantial parameter range. At extremely high parameter values, the system becomes chaotic (Fig. 5f), with windows of periodic (Fig. 5e) and quasiperiodic behavior. This transition is corroborated by the largest Lyapunov exponent of the closed curve as a function of *γ* (Fig. 5g).

**Fig 5:**
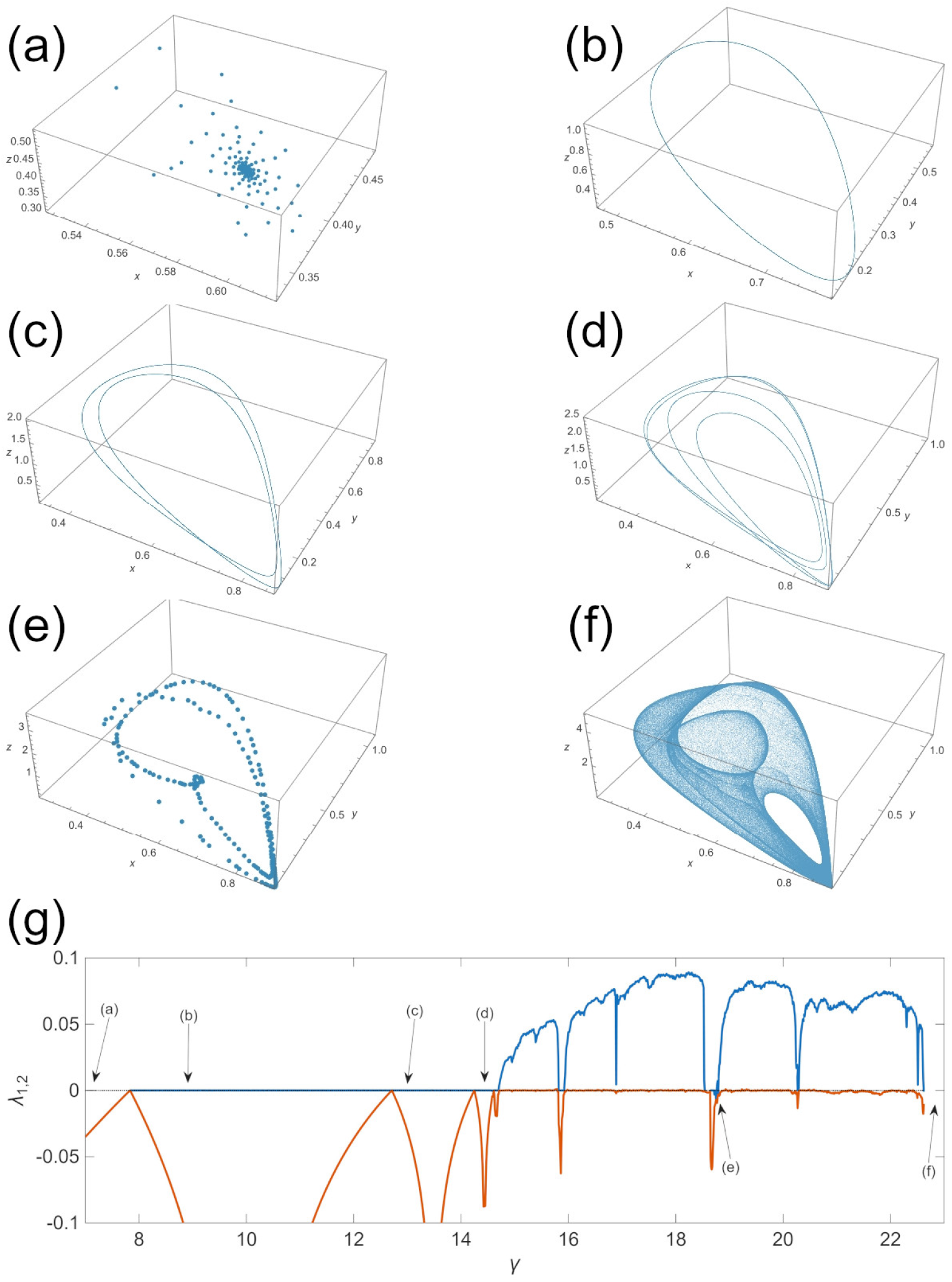
Dynamics of the system with Beverton–Holt host dynamics near the Neimark– Sacker bifurcation. (a)–(f): attractors obtained from numerical orbit calculations. (g): Lyapunov exponents as functions of *γ*; only the largest two exponents are shown. In all cases, the parameter values were fixed at *α* = 0.1 and *β* = 3.3. The value of *γ* was set at (a) *γ* = 7, (b) *γ* = 9, (c) *γ* = 13, (d) *γ* = 14.5, (e) *γ* = 18.75, (f)*γ* = 23.

In this case, rich dynamical behavior occurs only at extreme parameter values. Biologically, populations in this model are therefore more resilient than those with Ricker-type host fecundity, given the extremely low population levels that can arise from chaotic or extreme oscillatory behavior.

## 4 Discussion

In this work, we have defined and studied a model of hyperparasitism based on wasps of the family Trigonalidae. We assume that hosts (caterpillars) ingest hyperparasitoid eggs laid by trigonalid wasps and subsequently incubate a larva. If a host carrying a trigonalid larva is parasitized by a primary parasitoid, commonly another wasp species, the host and the primary parasitoid larva die, while the hyperparasitoid larva successfully develops into an adult. Otherwise, the host continues its life cycle normally and the hyperparasitoid larva dies. We consider two host fecundity functions in the absence of parasitism: Ricker- and Beverton–Holt-type functions. These functions correspond, respectively, to scenarios in which the primary parasitoid is the host’s main enemy and in which other enemies also regulate the host population.

By considering different host growth regimes, represented by high and low values of *α*, our model suggests that hyperparasitoids thrive when the host is prolific but fare worse when it reproduces slowly. Under Ricker-type host dynamics, the model exhibits mathematically rich behavior. However, the range of parameter values and initial conditions for which the hyperparasitoid survives and all three populations coexist without dangerous, extreme oscillations is very narrow, especially when the host is prolific. Indeed, a previous analysis of this model, which was restricted to delimiting the region of stability of the coexistence equilibrium in parameter space, concludes that […] if one asks how such a [host-parasitoid-hyperparasitoid] system could be constructed, it seems clear that much of this parameter space is unlikely to be realized by species in the real world; as the original host-primary relationship would have been too unstable to be likely to persist. [14]

In contrast, the model with Beverton–Holt-type host dynamics exhibits a smaller range of qualitative behaviors as the parameters change, while the hyperparasitoid persists across a much wider range of initial conditions and parameter values. This persistence represents resilience to both fluctuations in population size and environmental or behavioral changes. Furthermore, chaotic dynamics arise only at extremely high, and therefore mostly unrealistic, parameter values. Paradoxically, the absence of other host enemies promotes uncontrolled host growth, leaving hyperparasitoids susceptible to host population crashes even though both parasitoids and hyperparasitoids rely on abundant hosts for survival. Our minimal model therefore agrees with much more complicated models [9], in which populations at the highest trophic levels are more sensitive to environmental changes than populations at lower levels. Our results show that this instability may arise directly from perturbations in populations at the lowest trophic levels. They also resolve the apparent paradox raised in [14]: for a hyperparasitoid system such as the caterpillar–ichneumonid–trigonalid system to persist in the real world, other enemies must regulate the host even in the absence of parasitoids, thereby preventing dangerous oscillations at lower trophic levels.

From the definitions of the nondimensional parameters and populations, the ratio of hyperparasitoids to hosts is *Z*_*n*_*/X*_*n*_ = (*D/γ*)(*z*_*n*_*/x*_*n*_). Because approximately one trigonalid wasp emerges from each parasitized host carrying a trigonalid larva, *D ≈*1. Table 1 shows that both Beverton–Holt scenarios and the Ricker scenario with low host fecundity predict approximately one hyperparasitoid per hundred hosts. This prediction agrees with observed trigonalid hyperparasitism rates of approximately 0.01 or less [25]. In contrast, the hyperparasitism rates reported in [26] range from 0.16 to 0.47. This discrepancy most likely arises because the native caterpillars in that study were parasitized by an introduced generalist tachinid flies rather than wasps. Additionally, these flies do not rely exclusively on caterpillars and also parasitize sawflies, a possibility not included in our model. However, the reported rates are also close to the coexistence prediction for the Ricker scenario with a highly fecund host. We therefore cannot rule out a scenario in which primary parasitoids are the hosts’ main enemies. The observations could also come from an oscillatory population sampled near a maximum. To our knowledge, no time-resolved data on trigonalid population sizes are available. This absence highlights the need for time-resolved ecological data rather than isolated observations.

**Table 1:** Predicted ratios of hyperparasitoids to hosts for typical stable coexistence states.

| Host dynamics | Host fecundity | Parameter values | Coexistence equilibrium | Ratio of $Z_n$ to $X_n$ |
| --- | --- | --- | --- | --- |
| Ricker | Low | $\alpha = 0.85,$<br>$\beta = 2.7, \gamma = 6$ | $x^* = 0.49,$<br>$y^* = 0.44,$<br>$z^* = 0.07$ | 0.02 |
| Ricker | High | $\alpha = 4, \beta = 10,$<br>$\gamma = 4$ | $x^* = 0.73,$<br>$y^* = 1.08,$<br>$z^* = 1.50$ | 0.51 |
| Beverton–Holt | Low | $\alpha = 0.5,$<br>$\beta = 3.2, \gamma = 20$ | $x^* = 0.36,$<br>$y^* = 0.16,$<br>$z^* = 0.05$ | 0.007 |
| Beverton–Holt | High | $\alpha = 0.1, \beta = 2,$<br>$\gamma = 6$ | $x^* = 0.62,$<br>$y^* = 0.32,$<br>$z^* = 0.06$ | 0.02 |

Based on classic entomological literature [10, 13], we assume that hosts carrying a trigonalid larva develop normally in the absence of further parasitism. We therefore exclude facultative primary parasitoids such as *Taeniogonalos venatoria*. To the best of our knowledge, no experimental evidence indicates whether hosts carrying a trigonalid larva are more susceptible to primary parasitoids, for example because of altered behavior or delayed metamorphosis. If their susceptibility differs, the model would need to distinguish between hosts with and without a trigonalid larva and assign different parasitism probabilities to the two subpopulations. To model *T. venatoria*, the system would also need to account for host death and trigonalid reproduction when a host carrying a trigonalid larva escapes further parasitism. Finally, we assume that the probability that a host carries a trigonalid larva depends only on the number of trigonalid eggs, as determined by the trigonalid population. At high host densities, however, increased herbivory may increase the number of encounters between hosts and trigonalid eggs. This effect could be incorporated by modeling vegetation explicitly or by including host abundance in the encounter probability.

## Declarations

### Author contribution

ZGR: Formal Analysis, Visualization, Writing - original draft, Writing - review and editing JMNS: Conceptualization, Formal Analysis, Funding Acquisition, Investigation, Methodology, Supervision, Validation, Writing - orginal draft

### Data availability

Data sets obntained during the model’s numerical analysis are available from the corresponding author on reasonable request.

